# Genetic architecture of reproduction and longevity in historical Dutch cohorts

**DOI:** 10.64898/2026.06.02.729506

**Authors:** Richard Meitern, Peeter Hõrak

## Abstract

Life-history theory predicts genetic trade-offs between reproduction and survival, often invoked to explain ageing through antagonistic pleiotropy, but evidence in humans is mixed. We analysed a historical Dutch genealogical dataset (birth cohorts 1850–1915) to estimate pedigree-based heritabilities and genetic correlations among lifespan, offspring number, and ages at first and last birth (AFB and ALB). Bayesian animal models were fitted by sex in samples surviving beyond ages 45 and 13, with year of birth modelled by a natural spline. Lifespan heritability was approximately 0.19–0.23. Reproductive-trait heritabilities varied by sex and were generally higher in men. Cross-sex genetic correlations were substantial. Genetic correlations between offspring number and lifespan were weakly negative in women (-0.10), near zero in men aged 45+, and weakly positive in men aged 13+. AFB was positively genetically correlated with lifespan, especially in the 13+ sample. Offspring number was strongly genetically correlated with earlier AFB and later ALB, while AFB and ALB shared positive genetic covariance. Phenotypic associations changed little after adjustment for birth year, province, and urban/rural marriage location. Overall, results provide little support for a strong additive genetic fertility–longevity trade-off and instead highlight shared genetic influences on reproductive timing and viability.

## Introduction

Cost of reproduction is a central concept in the theory of life-history evolution and in evolutionary theories for ageing [1–3]. It refers to a life-history trade-off whereby greater investment in reproduction leads to reduced parental longevity and/or diminished capacity for future reproduction. Empirically detecting such trade-offs, however, is often challenging in observational studies because individuals differ in the total resources available for allocation to both reproduction and survival [4]. For instance, research in both pre-industrial and contemporary human populations has established that parental mortality risk is nonlinearly related to the offspring number: mortality risk declines as parity increases from zero to two or three children, but rises with further increases in reproductive output. Initial decline in mortality with increasing parity likely results from poorer overall health-status of nulliparous and low-parity individuals, as compared to parents of two and three children [reviewed in 5, 6].

The mechanisms underlying survival costs of reproduction in humans can be examined at multiple levels of analysis. The disposable soma theory [7] states that high investment in reproduction is obtained at the expense of resources that could be used for somatic maintenance and repair. Physiological and metabolic changes during pregnancy can have lasting effects on mothers, particularly with high parity, by increasing cardiovascular risk through weight gain, hyperlipidemia, insulin resistance, and related processes [8]. Accordingly, studies in developed countries link higher parity to greater prevalence or mortality from obesity, diabetes, and cardiovascular disease [9]. In developing countries, repeated pregnancies may lead to maternal depletion syndrome and long-term nutritional deterioration [10].

However, early proponents of the reproductive costs framework argued that reduced offspring production among long-lived women is grounded in evolutionary genetics, reflecting a heritable human life-history strategy characterized by a trade-off between fertility and longevity [11]. This perspective is rooted in the concept of antagonistic pleiotropy, whereby genetic variants that enhance some components of fitness simultaneously exert detrimental effects on others [12, 13].

Studies using quantitative-genetic, family-based and genomic approaches have yielded mixed evidence for genetic trade-offs between reproduction and survival. In preindustrial Finns, genetic correlations indicated shorter lifespans among women genetically predisposed to begin reproduction earlier or to have shorter interbirth intervals [14]. In the Framingham Heart Study, a strong negative genetic correlation between lifespan and offspring number was observed in women (r_g_= -0.88—-0.69) but none in men [15]. More recent genomic studies have also reported patterns consistent with antagonistic pleiotropy: genetic propensity for higher reproductive output was associated with increased cardiovascular disease risk [8, 16] and polygenic scores for earlier or greater reproduction predicted lower parental survival [17]. Recent meta-analysis found that majority of diseases that showed negative genetic correlations with longevity showed positive associations with fertility. The same study also observed a small negative genetic correlation between fertility and parental longevity (r_g_=-0.10) [18].

Other studies, however, have found weak, null or opposite relationships. A positive genetic correlation between the number of offspring surviving to age 20 and lifespan was detected in historical South Tyroleans [19]. In Swedish twins, the association between parenthood and longer survival was not explained by genetic or environmental factors shared between co-twins [20]. Quantitative-genetic analyses of a historical Utah population similarly argued against models of ageing that require a trade-off between reproduction and late-life survival [21]. Moreover, familial longevity was not associated with later reproduction or with a polygenic score for age at menopause in Dutch cohorts [22]. Mixed evidence is also found in non-human animals. A recent meta-analysis found an overall positive, rather than negative, genetic correlation between survival and other life-history traits, including fertility, although estimates were highly heterogeneous [23].

Here we use a large human genealogical dataset to test for a trade-off between reproduction and survival in a quantitative genetic framework. In historical human populations, number of ever born offspring is considered a relevant proxy of fitness [24, 25]. Offspring number is a composite trait that depends mechanically on the ages of reproductive onset and cessation and is genetically linked to both [14, 26–28]. We will therefore focus on detection of phenotypic associations and genetic correlations between lifespan, offspring number (parity) and ages of first and last reproduction (AFB and ALB).

We rely on an animal model, a generalized linear mixed-effects model that enables estimation of the additive genetic variances and genetic correlations between traits. It uses the information from all pedigree relationships to specify the expected phenotypic resemblance between relatives. The pedigree information was extracted from a crowd-sourced genealogical publicly available dataset [29]. The dataset is based on records of marriages conducted in the Netherlands between 1600 and 1999 and includes information on the births and deaths of spouses, their children, and their parents. The focal individuals included in our analyses were born between 1850 and 1915. To assess the evolutionary potential of lifespan and reproductive traits, we first estimated the heritabilities and cross-sex genetic correlations of these traits, and then calculated within-sex phenotypic and genetic correlations between reproductive traits and lifespan.

To reduce the problem of mechanical coupling between lifespan and reproductive opportunity, we restricted our main analysis to the individuals aged 45 and older. This age restriction reduces reverse temporal dependence: individuals dying before or during reproductive ages necessarily have reduced opportunity to reproduce. The 45+ analysis therefore focuses on individuals whose reproductive histories were largely observable before death. However, such conditioning on survival removes an important source of variation related to frailty and early-life viability [30, 31]. We therefore present the results for 13+ individuals in electronic supplementary material (ESM) as a complementary analysis showing how inclusion of early mortality changes the estimated associations.

## Methods

We analysed the Dutch publicly contributed genealogical marriage life-history dataset “Life histories of persons marrying, between 1600 and 1999, in the Netherlands, including children (GO924)” [29]. The dataset is based on records of marriages conducted in the Netherlands and includes information on the births and deaths of spouses, their children, and their parents. The focal individuals included in our analyses were born between 1850 and 1915.

During this period, the study population experienced a gradual improvement in biological standards of living. Adult stature increased across successive cohorts, indicating gains in net nutrition, health, and resistance to disease. These improvements coincided with rising real wages, declining mortality, and gradual changes in diet, hygiene, and living conditions, although substantial regional and social variation persisted. Childhood exposure to nutritional stress and infectious disease remained common, particularly prior to the full epidemiological transition, such that survival and reproductive outcomes were strongly shaped by environmental and socioeconomic conditions [32, 33]. Y-chromosome-based genealogical studies suggest that extra-pair paternity in the Low Countries averaged approximately 1.6% of births per generation over the past 500 years. Estimated rates varied substantially by population density and socioeconomic status, ranging from 0.4%–0.5% among farmers and middle- to upper-class families in sparsely populated communities to 5.9% among lower-socioeconomic-status families in the most densely populated cities. Rates in the latter group peaked in the late 19^th^ century [34].

Trait-specific samples are summarized in tables 1 and S1 in ESM. Phenotyped individuals were used for pairwise calculation of phenotypic correlations between traits. Their number exceeds that of individuals with relatives, which were used for genetic analyses. Each focal individual contributed a single per-person record. Offspring number was counted as the number of unique child identifiers pooled across all of that individual’s marriage records, so children from consecutive marriages are aggregated rather than undercounted. In this data subset the focal (married) individuals are all reproducers; parity is therefore conditioned on at least one child, and age at first and last birth (AFB, ALB) are defined only for reproducers. AFB and ALB were additionally missing for a small number of reproducers whose child birth years were not recorded (incomplete records rather than childlessness; about 2200 women and 2450 men in the 45+ sample). Data used for estimation of heritabilities of lifespan include also individuals with unknown offspring number, AFB and ALB.

Phenotypic associations between lifespan and reproductive traits were summarised as birth-year-adjusted partial correlations. For each sex and surviving-age sample, each trait in a pair was residualised on a natural spline of year of birth (ns(YOB, df = 5)) and the Pearson correlation between the residuals was calculated, so that both variables were adjusted for cohort trends. We also report the raw Pearson correlation and, as a test for regional sensitivity, a partial correlation and regression that additionally conditions on province (region) and urban vs rural location of marriage (ESM, table S2). Because associations between lifespan and reproductive traits were non-linear (figure 1 and figure S1 in ESM), we also report predicted lifespans for different parity categories (ESM, table S3) and group-specific linear slopes of lifespan on AFB and ALB (ESM, table S4).

**Figure 1.**
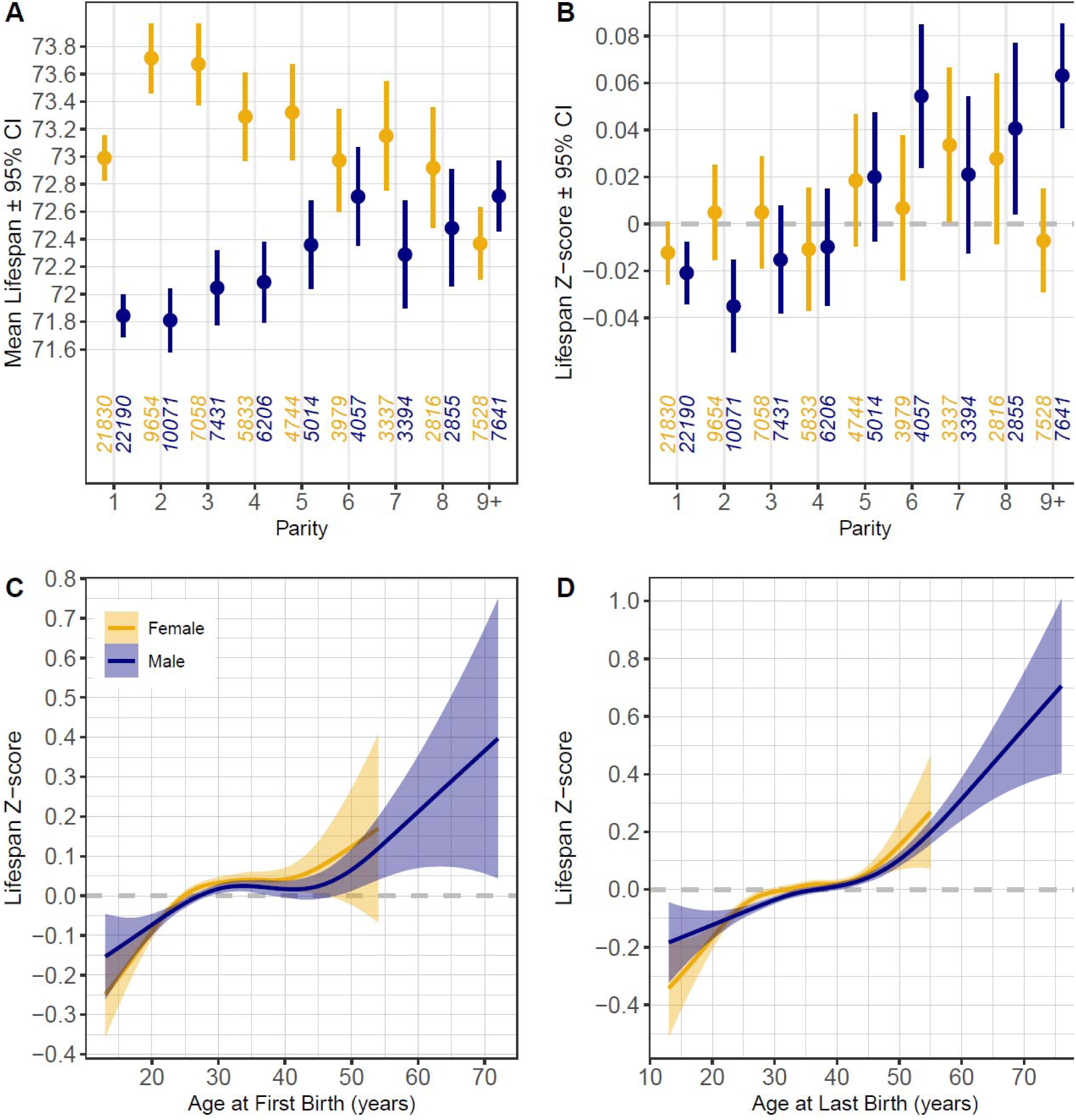
Associations between parity and crude lifespan (A) and lifespan z-scores, standardised within birth year and sex (B). Numbers at x-axis denote sample sizes per sex and parity. Birth-year adjusted predicted mean lifespans by parity category are shown in table S3 in the ESM. Associations between AFB (C) and ALB (D) vs lifespan z-scores, smoothed by GAM. Age is presented in years for the purpose of visualisation; age-group-specific linear slopes where both pairs of traits are adjusted for birth year are shown in the table S4 in ESM. Data for 45+ years old Dutch born between 1850 and 1915.

To assess the evolutionary potential of lifespan and reproductive traits, we estimated heritabilities and cross-sex genetic correlations for lifespan, number of children, AFB and ALB. We also estimated pairwise within-sex genetic correlations among these traits. Pedigrees were assembled from recorded parent–offspring links by adding missing parents as founders, removing invalid self-links, and sorting the pedigree for use in MCMCglmm.

Quantitative genetic parameters were estimated using Bayesian animal models implemented in MCMCglmm [35]. Three sets of models underlie the results reported here. Single-sex univariate models, fitted separately for women and men, provided the narrow-sense heritabilities of tables 1 and S1 in ESM. Cross-sex models treated the same trait measured in women and in men as two sex-limited responses and provided the cross-sex genetic and phenotypic correlations in those same tables. Within-sex bivariate models were fitted separately in each sex for pairwise combinations of lifespan, parity, AFB and ALB; these provided the genetic and phenotypic correlations among traits shown in figure 2 and figure S7 in ESM. Year of birth was entered as a trait-specific natural spline (ns(YOB, df = 5)) throughout.

**Figure 2.**
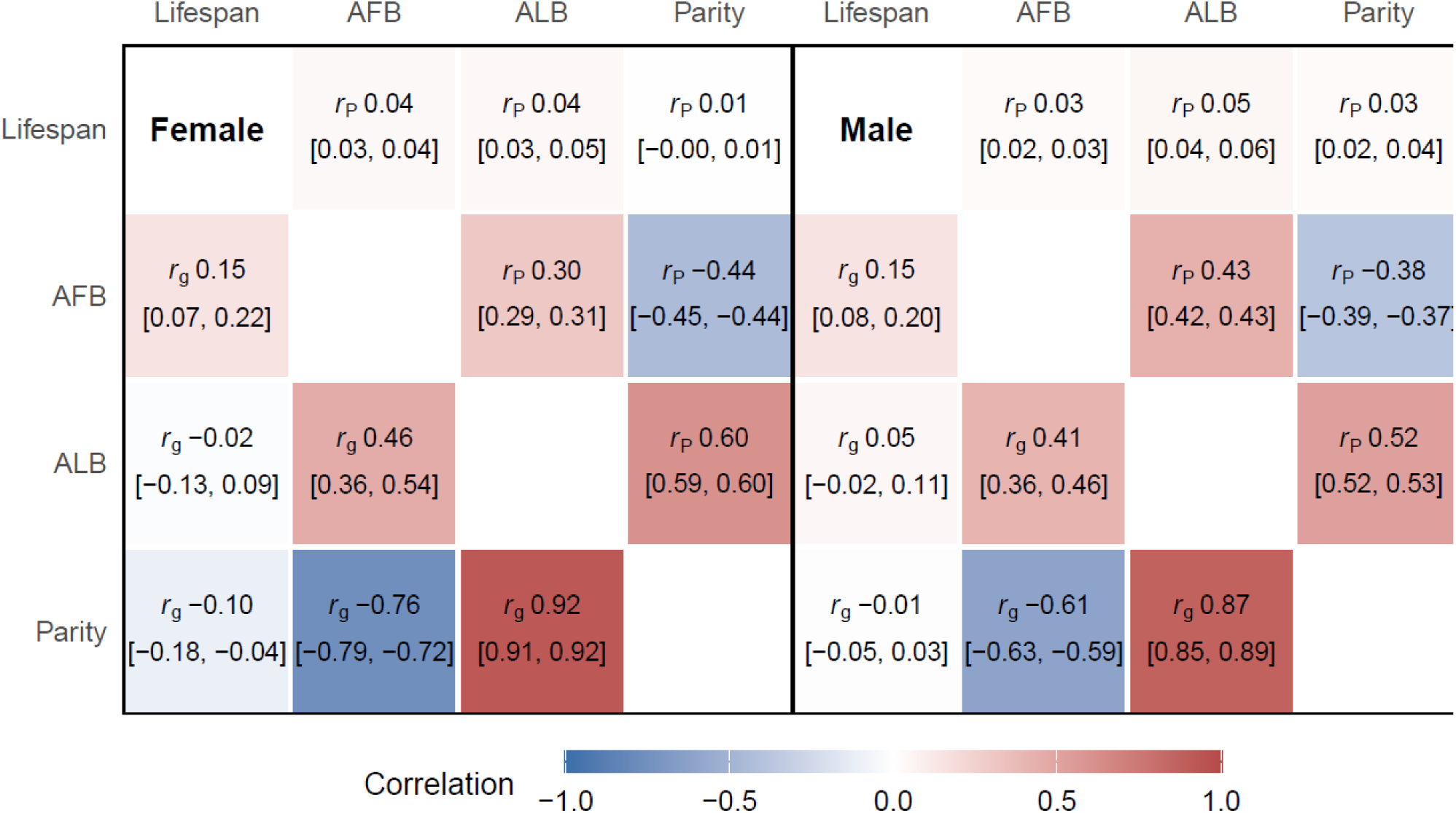
Phenotypic (above diagonal) and genetic (below diagonal) animal-model-derived correlations between lifespan and reproductive traits in 45+ sample. Numbers in parentheses denote 95% HPD intervals. Correlations with parity are converted to observed scale.

Lifespan, AFB and ALB were modelled as Gaussian traits with an identity link. Parity was modelled as a Poisson trait with a log link, so its additive genetic variance, heritability and genetic correlations are estimated on the latent log-rate scale, that is, the scale of the linear predictor for expected offspring number. On this scale they describe (co)variance in log expected fertility rather than in observed counts, and remain within their usual bounds (heritability in [0,1], correlations in [−1,1]). For interpretability we additionally report observed-scale heritabilities and correlations obtained at reference covariate values following [35]; genetic correlations involving parity are reported on the latent scale unless stated otherwise. An observation-level variance component was estimated for parity, absorbing extra-Poisson heterogeneity on the log scale (offspring counts were overdispersed relative to the Poisson expectation, with a variance-to-mean ratio of approximately 2.7 in both sexes and both age samples).

Additive-genetic and maternal-line effects were fitted with unstructured covariance matrices (us(trait):animal and us(trait):parent_line_id), so that the two additive-genetic variances and their covariance were estimated, with the genetic covariance identified from cross-trait resemblance among pedigree relatives. The maternal-line term groups individuals by mother identity and therefore captures resemblance among maternal sibships beyond that expected from additive genetic relatedness; individuals with no recorded mother were each assigned a unique level so that they were not pooled. The residual structure differed among model types. In within-sex bivariate models of two Gaussian traits (lifespan–AFB, lifespan–ALB, AFB–ALB) an unstructured residual matrix was fitted (us(trait):units), so that the individual-level residual covariance between traits was estimated. In cross-sex models the residual matrix was necessarily diagonal (idh(trait):units), because the two responses are sex-limited and are never observed in the same individual, leaving their residual covariance unidentifiable. In mixed Poisson–Gaussian models the residual matrix was likewise diagonal: the Gaussian residual variance and the Poisson observation-level variance were estimated, while the cross-trait residual covariance was fixed at zero for identifiability, so that the additive-genetic covariance is identified under the assumption of no observation-level cross-trait covariance.

Variance components were assigned inverse-Wishart priors. For Gaussian models these were weakly informative, with an identity scale matrix and degrees of freedom equal to the dimension of the (co)variance structure (V = I_p, nu = p; V = 1, nu = 1 in univariate models). For models including parity, mildly regularising priors were used on the latent log scale to improve mixing: for the additive-genetic and maternal-line structures V = 0.02 I_p (0.2 I_p in the cross-sex parity model) with nu = p + 2, and for the residual structure V = diag(0.5, 0.02) (0.5 I_p in the cross-sex parity model) with nu = p + 2. Parameter expansion was not used.

Chains were run with settings fixed a priori: a 30 000-iteration burn-in, no thinning, and one of two target run lengths. Long runs were executed as checkpointed batches concatenated into a single posterior sample, with the burn-in discarded once in the first batch and subsequent batches continuing the chain from the previous end state; because batches were of fixed size, realised chain lengths differ slightly from their nominal targets. Of the 48 models whose estimates are reported here, 18 were run to approximately 0.5 million iterations (470 000–490 000 retained samples) and 30 to approximately 1.3 million iterations (1 230 000–1 270 000 retained samples). Realised settings for every reported model are given in ESM, table S5.

Convergence and sampling efficiency were assessed per model on the variance–covariance samples using the Heidelberger–Welch stationarity test, the Geweke diagnostic, and effective sample sizes (coda). All variance–covariance components passed the Heidelberger–Welch stationarity test in 35 of the 48 models; Geweke pass rates were more variable and are reported per model in ESM, table S5. Median per-model effective sample sizes ranged from 273 to 4931. Effective sample size tracked run length rather than model type: every one of the 30 longer chains reached a minimum effective sample size above 200, whereas all 14 models with a minimum below 200 were among the 18 shorter chains. In particular, all 16 univariate models providing the heritabilities of tables 1 and S1 had minimum effective sample sizes of at least 232 (medians 759–3026). Posterior summaries are reported as posterior means with 95% highest-posterior-density (HPD) intervals; posterior means and medians were essentially identical for the reported quantities (maximum absolute difference 0.002 for heritability and 0.041 for a genetic correlation).

Analyses in the main text were conducted on individuals surviving to at least 45 years of age, by which time reproduction is largely complete (ESM, table S6). Parallel analyses of the full population aged above 13 are reported in the ESM.

## Results

### Demographic background and secular trends

Mortality was higher in women than in men between ages 25 and 45, whereas the pattern reversed after age 82 (ESM, Fig S1A). Among women, lifespan increased rapidly for cohorts born after approximately 1875–1880 (ESM, Figs S2B and S3B). This pattern was similar among women surviving to at least age 13 or 45. In contrast, among men who survived to age 45, mean lifespan fluctuated between 71 and 72 years for cohorts born between 1850 and 1895, and subsequently increased to nearly 74 years by the 1910 cohort (ESM, figure S2B). Among men in 13+ age group, increases in lifespan over the study period were minimal (ESM, figure S3B).

In both age-restricted subsets and sexes, parity declined gradually beginning with cohorts born around 1860 and levelled off among cohorts born after 1900 (ESM, Figs S2A, S3A). Ages at first and last reproduction showed similar U-shaped patterns across the study period in both age-restricted subsets, with broadly parallel trends in men and women (ESM, Figs S2CG, S3CD).

### Phenotypic associations between lifespan and reproductive traits

Among the 45+ women, lifespan peaked at parities of two to three children and declined at higher parities (Fig 1A). This pattern, however, faded when lifespans were adjusted for birth year averages, after which no clear-cut associations between offspring number and mothers’ lifespan could be detected (figure 1B, ESM, figure S4). Among women in 13+ age group lifespan showed positive associations with parity which were similar on raw and birth-year adjusted data (r = 0.06 and 0.08; ESM, Figs S5, S6AB).

Among men, lifespan increased with parity irrespective of whether raw or adjusted data were analysed. In the 13+ age group this relationship was nearly monotonic and markedly stronger in the 13+ age group than in the 45+ age group (r = 0.09 vs 0.03, respectively; figure 1AB, figure 2, ESM, Fig S4, S5, S6AB).

In the 45+ sample, AFB was weakly and positively correlated with lifespan in both sexes (Fig 1CD, ESM, Figs S4, S5, S6CD). This positive correlation was stronger among individuals (particularly women) who started and ended their reproduction before age 25 and after 40. Among the bulk of population who started and ended their reproduction between ages 25 and 40, lifespan was virtually independent of AFB and ALB (ESM, table S4). In the 13+ dataset, the associations between reproductive timing and lifespan differed from those in the 45+ sample. Lifespan increased nearly monotonically with AFB and ALB among both men and women (SI Appendix, Figs S4, S5, S6CD).

Correlations between AFB and ALB were moderate and in a similar magnitude in both age groups and when calculated on the basis of raw data and z-scores (Figs S4, S5). Correlations for men (r = 0.42 – 0.44) were stronger than correlations for women (r = 0.30 – 0.34).

### Heritabilities and genetic correlations

Among individuals who had reached age 45, heritability of lifespan was 0.21 for women and 0.19 for men (table 1), which was similar to heritabilities calculated on the basis of the whole sample (0.23 for both sexes; ESM, table S1). Heritability of offspring number was substantially higher in men (h^2^_obs_ = 0.44 and 0.43 in the 45+ and 13+ samples) than in women (h^2^_obs_ = 0.17 and 0.15; table 1, ESM, table S1). Heritabilities of AFB (h^2^ = 0.34 – 0.49) and ALB (h^2^ = 0.13 – 0.35) were higher in men than in women (table 1, ESM, table S1).

**Table 1.**
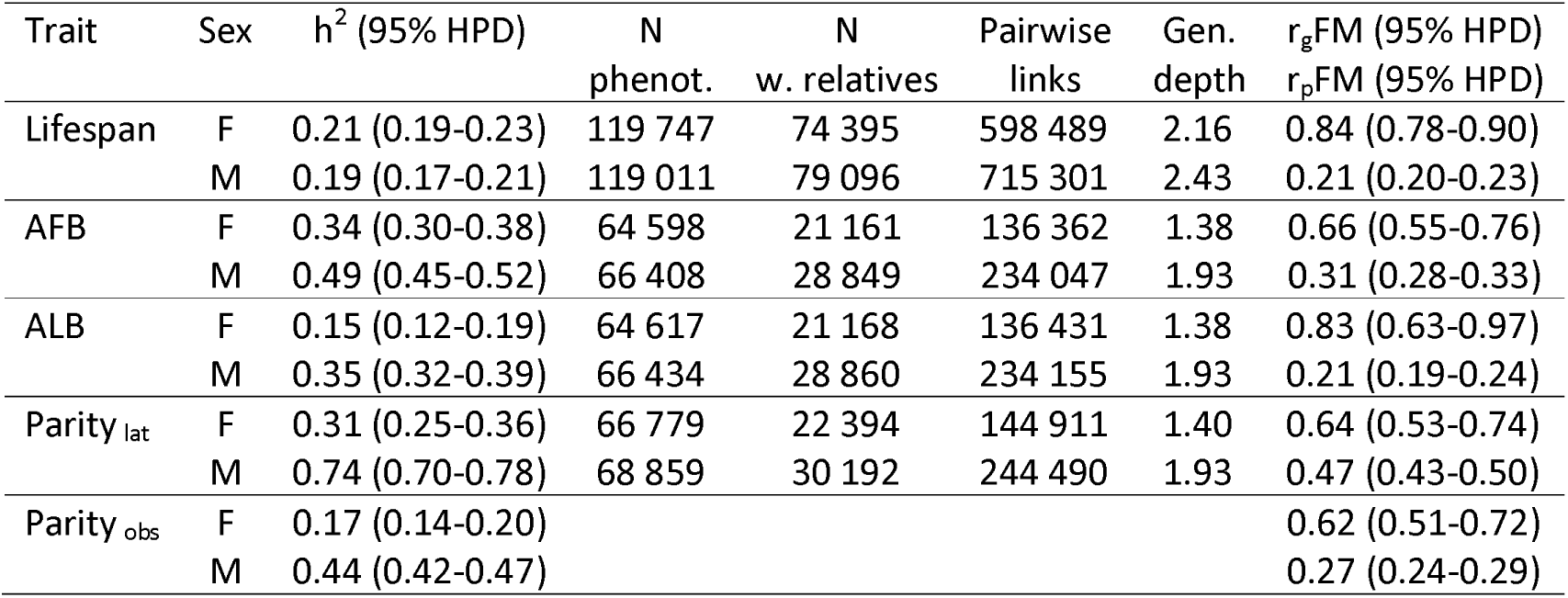
Heritabilities for lifespan and reproductive traits among individuals who had reached age 45. Heritabilities are from single-sex univariate animal models; r_g_FM and r_p_FM are cross-sex genetic and phenotypic correlations from bivariate models in which the same trait in women and men was treated as two sex-limited responses. N phenot. is the number of phenotyped individuals for each trait. N w. relatives is the number of phenotyped individuals having at least one phenotyped parent, offspring or sibling. Pairwise links counts relative pairs with r ≥ 0.125 (first cousins and closer); the model uses the complete pedigree, including more distant relationships. Gen. depth is average ancestral generation depth (maximum is 14). Narrow sense heritability estimates (h^2^) are presented as posterior means with the associated 95% Highest Posterior Density (HPD) intervals on the latent scale (h^2^ _lat_) and, for parity also converted to the observed scale (h^2^_obs_) [35]. h^2^_obs_ was calculated at reference values of 3.1 children for females and 3.0 children for males. r_g_FM and r_p_FM are cross-sex genetic and phenotypic correlations between trait values.

Cross-sex genetic correlations indicated substantial shared genetic architecture between sexes. These correlations were especially high for lifespan and ALB (0.84 and 0.83) followed by estimates for AFB (0.66) and parity (0.64) in 45+ age group (table 1). In 13+ age group these correlations were similar in magnitude, ESM, table S1).

Among women, genetic correlations between lifespan and offspring number were weakly negative (-0.10) and identical in 13+ and 45+ samples (figure 2, ESM, figure S7). Among men in 45+ sample, genetic correlation between lifespan and parity did not differ from zero while in men aged 13+ years, the correlation was weakly positive (0.05). Genetic correlations between ALB and lifespan were negative (-0.11) among 13+ women and indistinguishable from zero in three other age-sex groups. AFB correlated with lifespan positively and most strongly so among 13+ women (r_g_ = 0.40) and men (r_g_ = 0.36) while in the 45+ sample the correlation was 0.15 among both sexes.

Parity correlated strongly and negatively with AFB in both sexes and age groups (r_g_ = -0.59 to -0.76) while parity’s correlations with ALB were strong and positive (r_g_ = 0.87 – 0.92; figure 2; ESM, figure S7).

## Discussion

The cost of reproduction is a cornerstone of life-history theory and evolutionary explanations of ageing, predicting that greater reproductive investment should come at the expense of survival and longevity [1–3]. In humans, this idea has long been invoked to explain negative associations between reproductive traits and lifespan observed in historical and contemporary populations [17, 36, 37]. However, distinguishing between physiological costs, environmental heterogeneity, and genuinely genetic trade-offs has proven difficult [38]. Using a large multigenerational genealogical dataset and a quantitative genetic animal-model framework, we test whether reproduction and survival are linked by antagonistic genetic architecture. Lifespan was moderately heritable in both sexes, and reproductive traits—especially offspring number and reproductive timing in men—also showed appreciable heritability. Despite weak and sometimes inconsistent phenotypic associations between reproductive traits and survival, we found some evidence for negative genetic correlations between reproductive traits and lifespan in women and positive genetic correlations between AFB and lifespan in both sexes. Yet the genetic correlations between parity and lifespan were small: weakly negative in women, null or weakly positive in men. Thus, our results are broadly consistent with the life-history prediction that human longevity may be constrained by genetic trade-offs with reproduction, but they do not support a strong additive genetic fertility–longevity trade-off.

### Secular trends and phenotypic associations

The cohorts examined here span a period of rapid demographic and epidemiological transition in the Netherlands, characterized by declining fertility and increasing longevity. These secular trends closely mirror patterns observed across much of Europe during industrialization and reflect profound changes in nutrition, disease exposure, and living conditions [39, 40]. Against this background, we failed to identify the common twentieth-century inverse J-shaped pattern, in which parental survival peaked at intermediate parities and declined as the offspring number increased further [5, 6, 41]. Among men, lifespan increased modestly with parity, with no indication of reduced survival at high reproductive output. The absence of increased male mortality at high parities differs from findings of previous studies. One possible reason might be over-representation of older elite men in genealogical samples relative to the general population, partly because long-lived male lineages are more visible and more likely to have surviving descendants who contribute their trees [42, 43].

On the other hand, phenotypic correlations between other reproductive traits and parental longevity were in the same direction as those reported in other pre-industrial and contemporary human populations [reviewed by 44, 45, 46]. The positive correlation between lifespan and ALB (r = 0.24) was stronger than the correlation between lifespan and AFB (r = 0.13) in the full 13+ sample. A proximate explanation for the association between late reproduction and long lifespan is that individuals with a propensity for long life possess better physiological condition, health, and/or material and social resources already during reproductive age, and can, therefore, afford to reproduce in older age than potentially short-lived persons who lack such resources [see 4, 38, 46].

### Genetic architecture of lifespan and reproduction

We found moderate and relatively stable heritability of lifespan (0.19–0.23) across sexes and age-restricted samples, with high cross-sex genetic correlations. Our lifespan heritability estimates align well with previous quantitative genetic studies in historical populations [14, 19, 47–49], supporting the view that lifespan retains substantial evolutionary potential despite strong environmental influences. Our cross-sex genetic correlations of lifespan were remarkably similar to those recorded in 19^th^ century Utah (r_g_ = 0.82) [21], suggesting that lifespan in men and women is influenced largely by shared genetic factors [see also 50].

In contrast, parity showed pronounced sex differences in heritability, whereas latent-scale heritability estimates for parity were relatively high (0.73—0.74) in men. Our latent-scale estimates are not directly comparable to published records that have treated parity as a Gaussian trait [35]. However, when converted to observed scale, our estimates of heritability for offspring number were comparable to previously published findings [14, 33, 51, 52]. However, contrary to the findings of this study, offspring number was substantially more heritable in women than in men in a pedigree based study of Finns born between 1745 and 1900 [14]. The authors suggest that male fitness in their study population might have depended mainly on the reproductive quality of their spouses. It is possible that in the current study of Dutch cohorts born between 1850 and 1915, the heritable effects of female phenotypic quality on their offspring number were less pronounced than the heritable effects of male quality (including resources).

Genetic correlation between parity and lifespan was indistinguishable from zero among 45+ men and weakly positive in the 13+ sample (r_g_ = 0.05). These results differ from the findings from 19^th^ century Utah [21] where male—but not female—post-50 lifespan correlated genetically positively with fitness (r_g_=0.11). In our study, the genetic correlation was -0.10 for both age groups in women. This estimate exactly matches the genetic correlation of -0.10 between fertility and parental longevity reported in a pooled-sex GWAS meta-analysis [18]. These results contrast with findings of the Framingham Heart Study, which reported a strong negative genetic correlation (r_g_ = -0.88—-0.69) between lifespan and offspring number in women [15]. However, that study relied on a modest sample of 5133 women, and the significance of genetic correlation depended on partitioning of the covariance structure.

As with parity (and possibly for the same reasons), heritability estimates for both AFB and ALB were substantially higher in men than in women. These estimates were of similar magnitude to those reported in previous studies [reviewed by 27, 53]. Cross-sex genetic correlations were again relatively high (0.66—0.83). The strong genetic correlations of AFB and ALB vs parity were remarkably similar between the sexes suggesting little evidence for genetic sexual conflict. This result is consistent with findings in a historical Finnish population [27].

Positive genetic correlations between AFB and lifespan are consistent with those found in a meta-analysis of European-origin populations [54] and also with findings of GWAS studies in UK biobank showing that genetic variants associated with later age at first birth are associated with a longer life span of mothers [55] or both parents [17, 56] of participants. The study by Long and Zhang [17] provides evidence for the horizontal antagonistic pleiotropy between reproduction and life span as the negative genetic correlations involving reproductive traits were at least partly independent of realized reproductive output. In the current study, the positive additive genetic correlations between AFB and lifespan are consistent with shared genetic influences linking later reproductive onset to longer lifespan. Their greater magnitude relative to the phenotypic correlations (figure 2; ESM, figure S7) is compatible with dilution by non-genetic variance or countervailing environmental covariance, although the present comparison does not by itself demonstrate environmental masking or antagonistic pleiotropy.

Genetic correlation between parity and AFB ranged from -0.59 to -0.76 across sexes and age subsets, corroborating findings of pedigree [27] and GWAS studies, that early reproduction is genetically positively linked to reproductive success [55, 57–60]. Genetic correlation between parity and ALB were even stronger in magnitude (r_g_ = 0.87–0.92). These estimates were again in the similar magnitude with the correlations detected in a historical Finnish population [27]. Interestingly, a single GWAS study that has assessed genetic correlation between ALB and offspring number found a negative relationship (r_g_= -0.40) while the phenotypic correlation between these traits was positive (r =0.22) [58].

Genetic correlations between AFB and ALB were largely similar and relatively strong across sexes and age subsets (r_g_= 0.41—0.56). These correlations provide evidence for a genetic constraint on achieving the combination of early onset and late cessation of reproduction. Previously, even stronger genetic correlation between AFB and ALB (r_g_= 0.91) was found in a small-scale genealogical study in 19^th^ century Swiss men [61] and in a GWAS study of female participants in UK biobank (r_g_ = 0.94) [58].

### Limitations and implications

A common concern when using genealogical data to study survival and reproduction is that such datasets are inherently selective, over-representing individuals who survived to reproduce and whose lineages persisted over generations. Early deaths, childless individuals, and socially marginal individuals — particularly men — are likely to be under-represented [42, 43]. Selection into the analysed sample can bias both phenotypic and genetic covariance estimates when inclusion depends on the traits under study or on unobserved causes of those traits. In our case, individuals who died before reproduction and individuals without recorded descendants are under-represented, so estimates should be interpreted as applying primarily to individuals who survived to the relevant age and entered the genealogical record. In this Dutch dataset, the number of phenotyped men and women was fairly similar for each trait, but the number of individuals with registered relatives and pairwise links was substantially higher for men than women (table 1, tables S1 in ESM). Pedigrees of men thus appear, on average, more complete than those of women.

Further, animal models estimate additive genetic covariance using pedigree resemblance, but if the pedigree is socially or geographically structured, relatives may resemble one another not only genetically but also because they share class, region, religion, cultural fertility norms, migration patterns, or record quality. If such clustering is not modelled, some “genetic” covariance may absorb shared environmental or population-structure covariance [62]. Unfortunately, the analysed dataset lacks the information about social class or religion of the participants. However, we tested for regional sensitivity by calculating phenotypic partial correlations between life-history traits that additionally conditions on province (region) and urban vs rural location of marriage (ESM, table S2). Partial correlations were largely unchanged after additional adjustment for province and urban/rural marriage location (max |Delta r|= 0.011), providing little evidence that these measured geographic factors materially confound the associations beyond birth cohort. This does not exclude residual confounding by other socioeconomic characteristics. However, if non-genetic/cultural inheritance inflated our estimates of heritability of reproductive traits, then such effect would be sex-specific as heritabilities of offspring number, AFB and ALB were all higher for men than women (table 1, table S1 in ESM).

Overall, this study confirmed that reproductive success is a composite trait shaped mechanically by the ages at reproductive onset and cessation, and genetically linked to both through shared genetic covariances. Despite detection of only weak negative genetic correlations between offspring number and maternal lifespan (and weakly positive or null relationship in men), genetic correlations between AFB and lifespan were moderate and positive, particularly in the full sample. These results suggest that shared genetic influences on viability, developmental tempo, and reproductive scheduling may be more important for manifestation of survival cost of reproduction than a simple allocation trade-off between reproduction and somatic maintenance.

The strong genetic correlations between parity and reproductive timing also emphasize that offspring number is not an isolated trait. Genetically, higher parity was associated with earlier first reproduction and later last reproduction, particularly through the extension of the reproductive span. Consequently, any association between parity and lifespan may partly reflect the genetic and demographic architecture of reproductive timing. This is important for interpreting offspring number as a proxy for fitness: although parity captures realized reproductive success, it is also conditioned by marriage, survival, fecundity, social norms, and cohort-specific fertility decline. Such interpretation is also relevant in epidemiological context: two studies on health and reproduction across 11 – 13 European countries found that birth timing was more relevant than parity for health outcomes in later life [63, 64].

Another important life-history trade-off emerging in this study is indicated by the relatively strong and consistent positive genetic correlation between AFB and ALB. In the context of very strong and opposite-sign genetic correlation between AFB and ALB vs parity, a positive genetic correlation between AFB and ALB could constrain an evolutionary response to selection on both traits [61]. This genetic trade-off may well contribute to the maintenance of within-population genetic diversity of reproductive timing which is an integral part of the slow/fast pace-of-life-syndrome (POLS) hypothesis [see 65, 66, 67].

Finally, findings from this historical Dutch population provide a useful point of comparison with GWAS studies in modern populations. GWAS and genealogical studies are based on different birth cohorts, exposed to different epidemiological, nutritional, and social conditions, and subject to different forms of sample-selection bias. They also rely on different analytical approaches for estimating genetic correlations. Nevertheless, several correlations observed here—including the genetic association between later reproductive onset and longer lifespan, the strong genetic link between earlier reproduction and higher parity, and the weak negative association between parity and lifespan—were similar in direction and magnitude to estimates from GWAS studies of contemporary populations. Strong cross-sex genetic correlations for reproductive traits and lifespan have also been observed in both types of studies. These similarities suggest that some components of the genetic architecture underlying human reproductive timing, parity, and lifespan have remained broadly stable across recent historical periods.

## Data, Materials, and Software Availability

The original data, along with the R code, are available at https://doi.org/10.5281/zenodo.18549502

## ACKNOWLEDGEMENTS

The study was funded by the Estonian Research Council grant PRG1137.

