## Supplementary Information file: Tables S1-S6 and figures S1-S7 for "Genetic architecture of reproduction and longevity in historical Dutch cohorts"

**Electronic Supplementary Material**

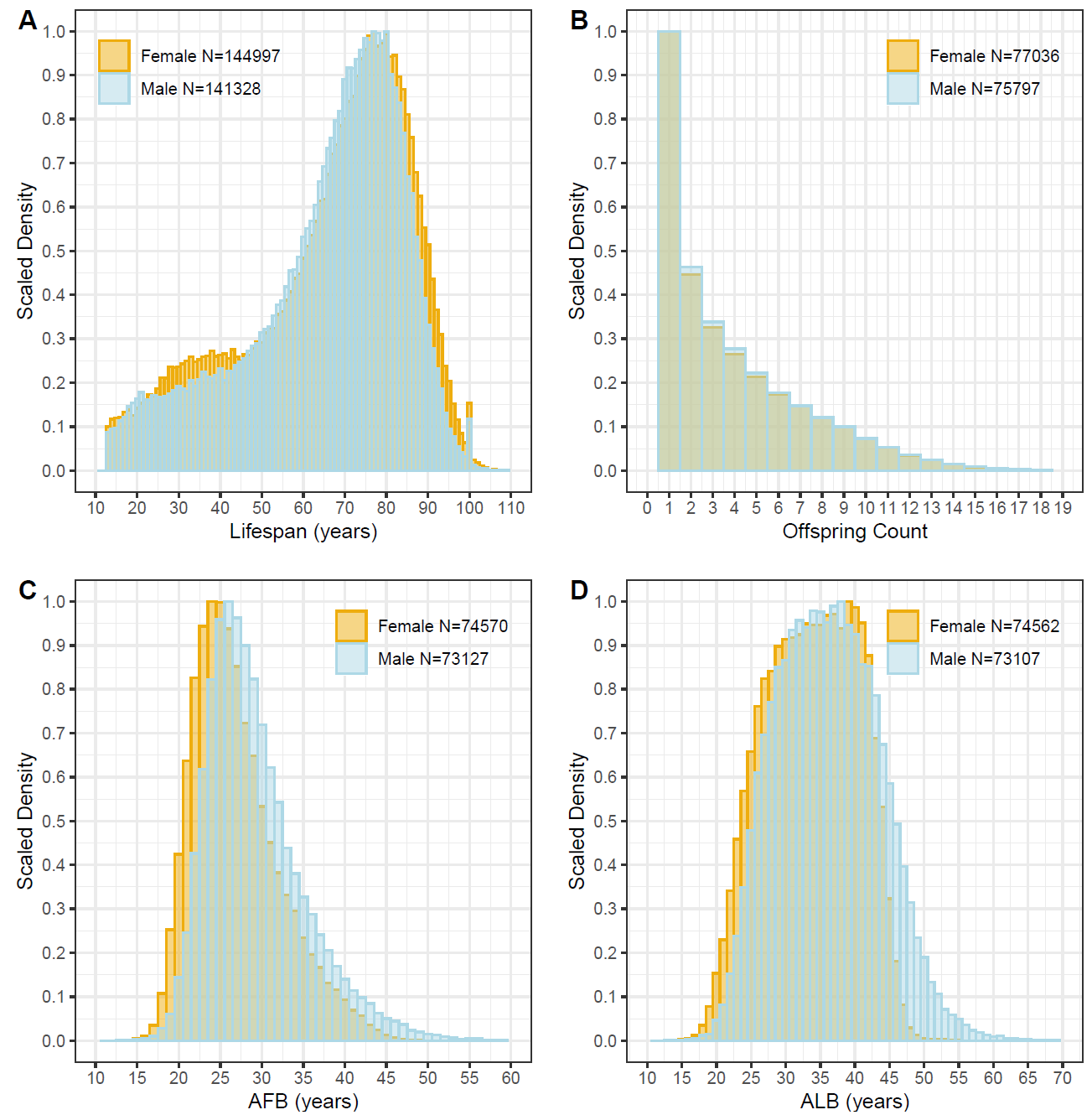

**Figure S1.** Distributions of lifespan (A), parity (B), age of first birth (C) and age of last birth (D) among 13+ years old Dutch born between 1850 and 1915.

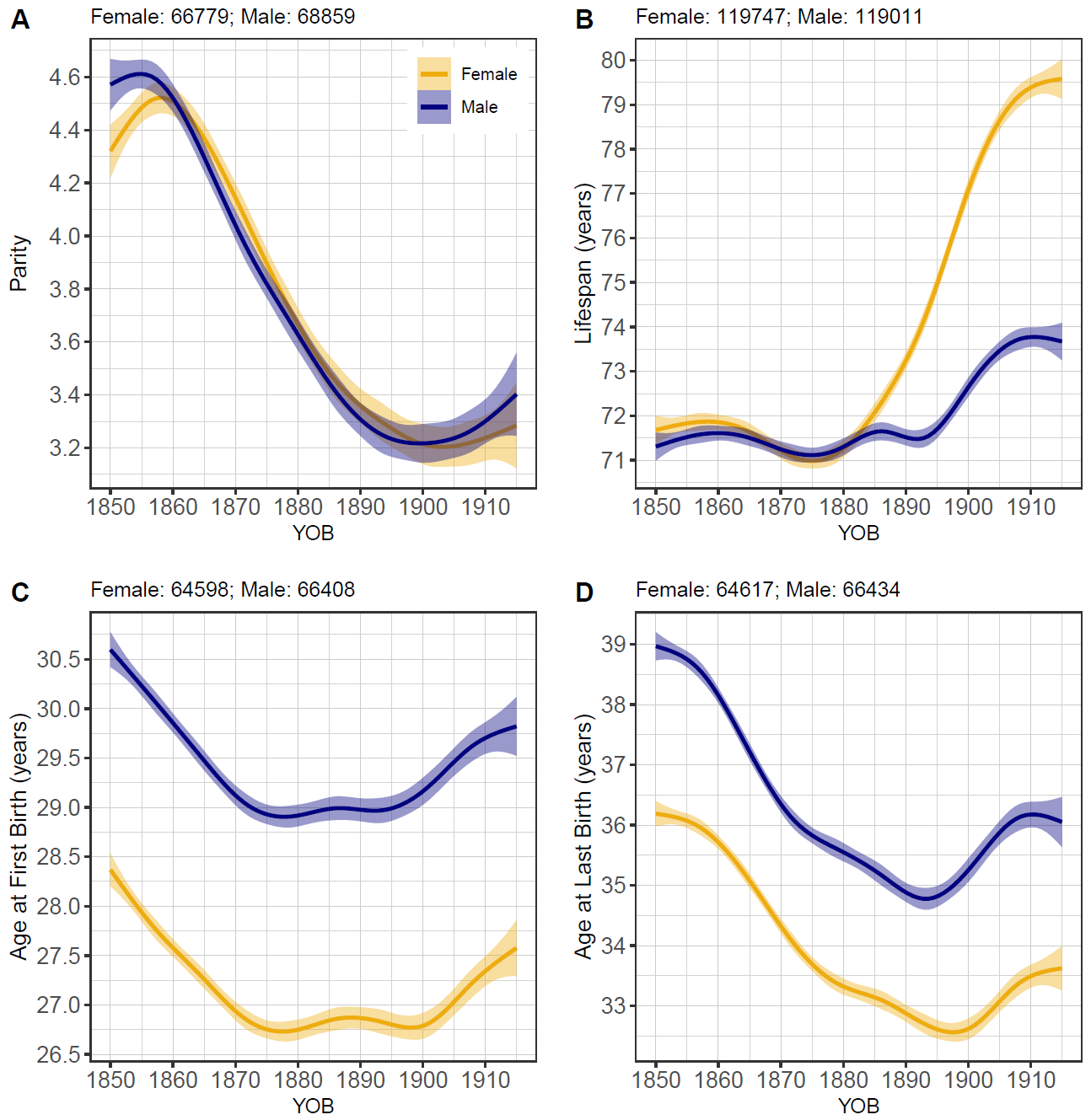

**Figure S2.** Secular trends in lifespan and reproductive traits in 45+ sample.

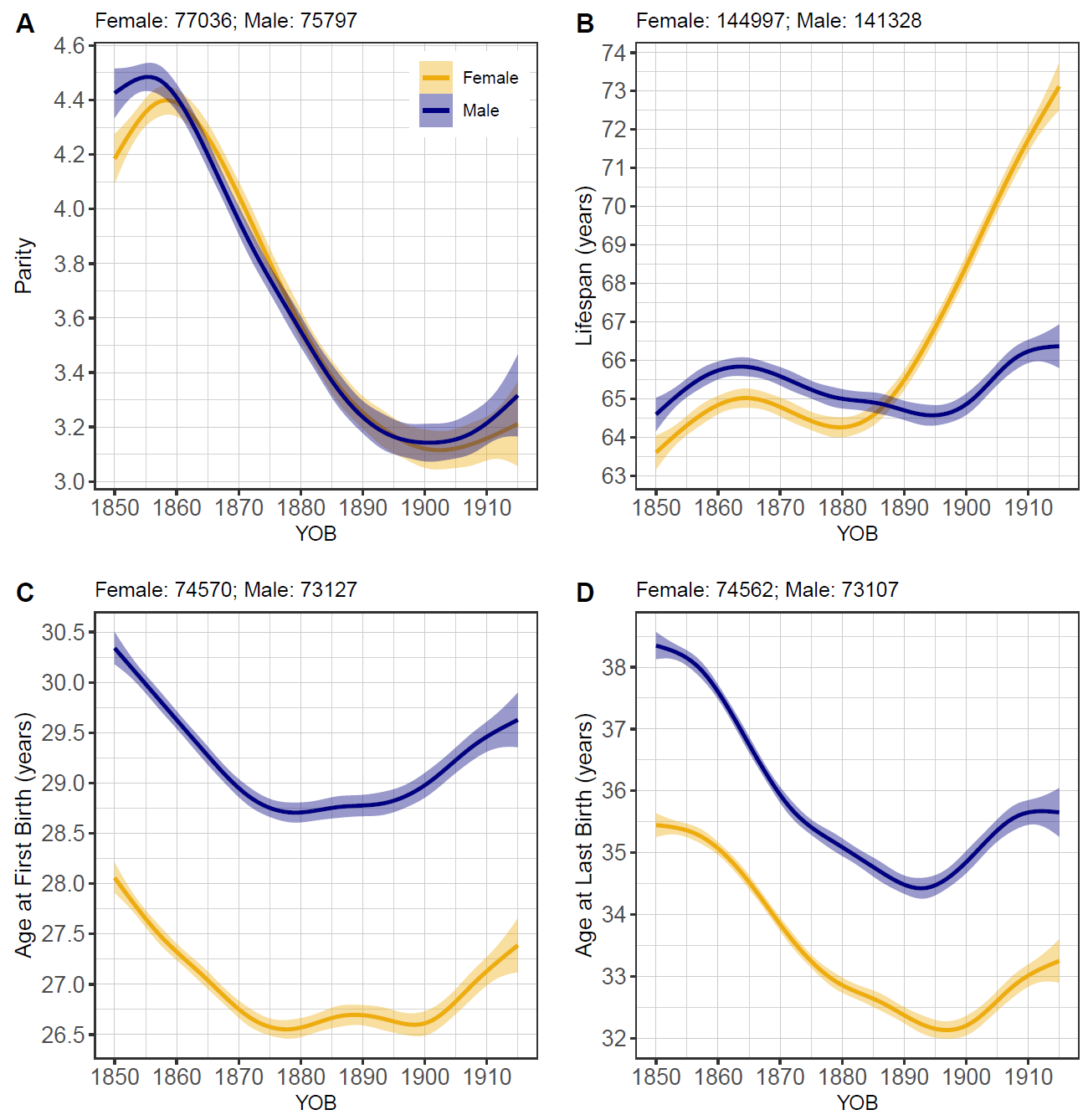

**Figure S3.** Secular trends in lifespan and reproductive traits in 13+ sample.

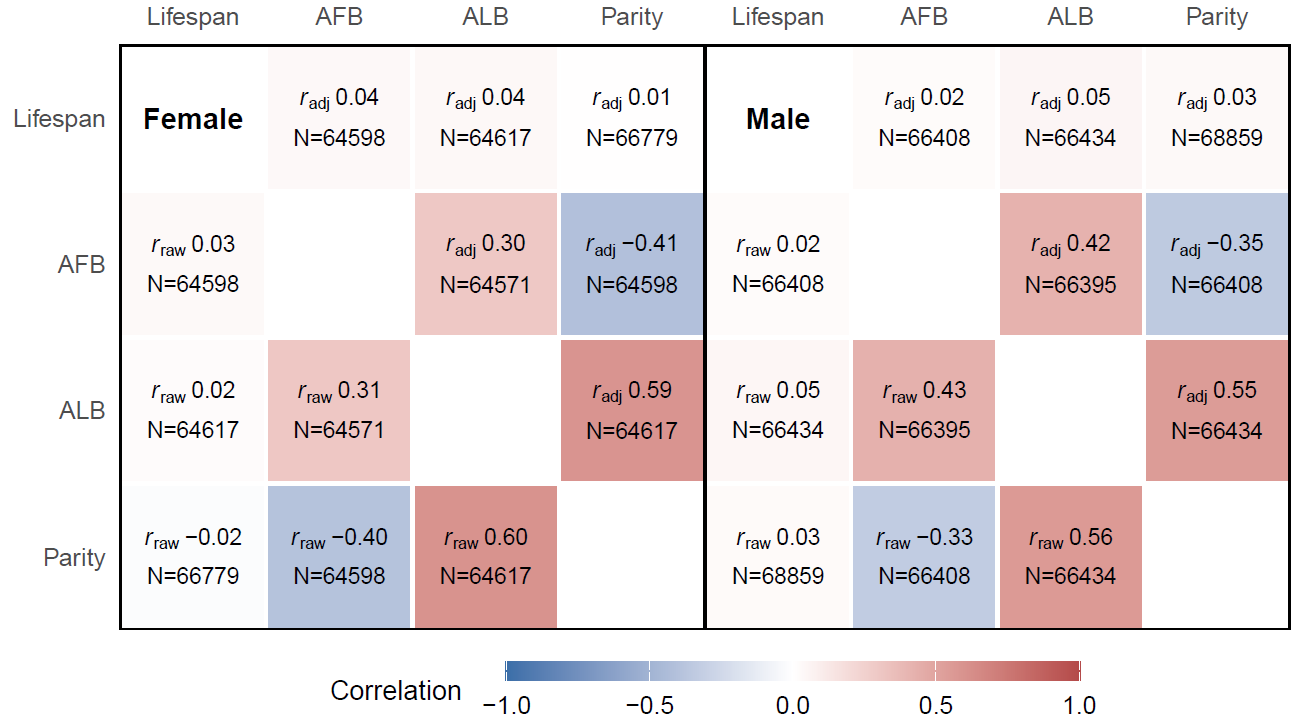

**Figure S4.** Raw (below diagonal) and birth-year-adjusted correlations between lifespan and reproductive traits in 45+ sample, calculated on the basis of phenotypic values.

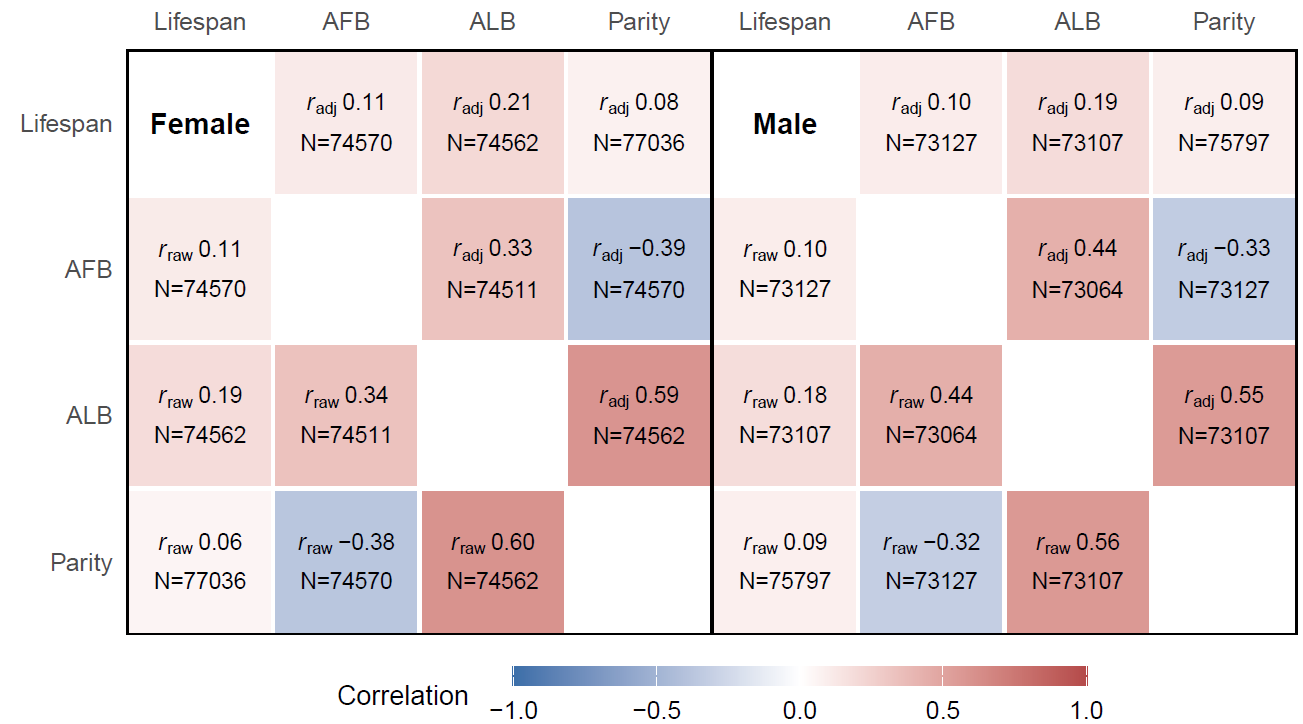

**Figure S5.** Raw (below diagonal) and birth-year-adjusted correlations between lifespan and reproductive traits in 13+ sample calculated on the basis of phenotypic values.

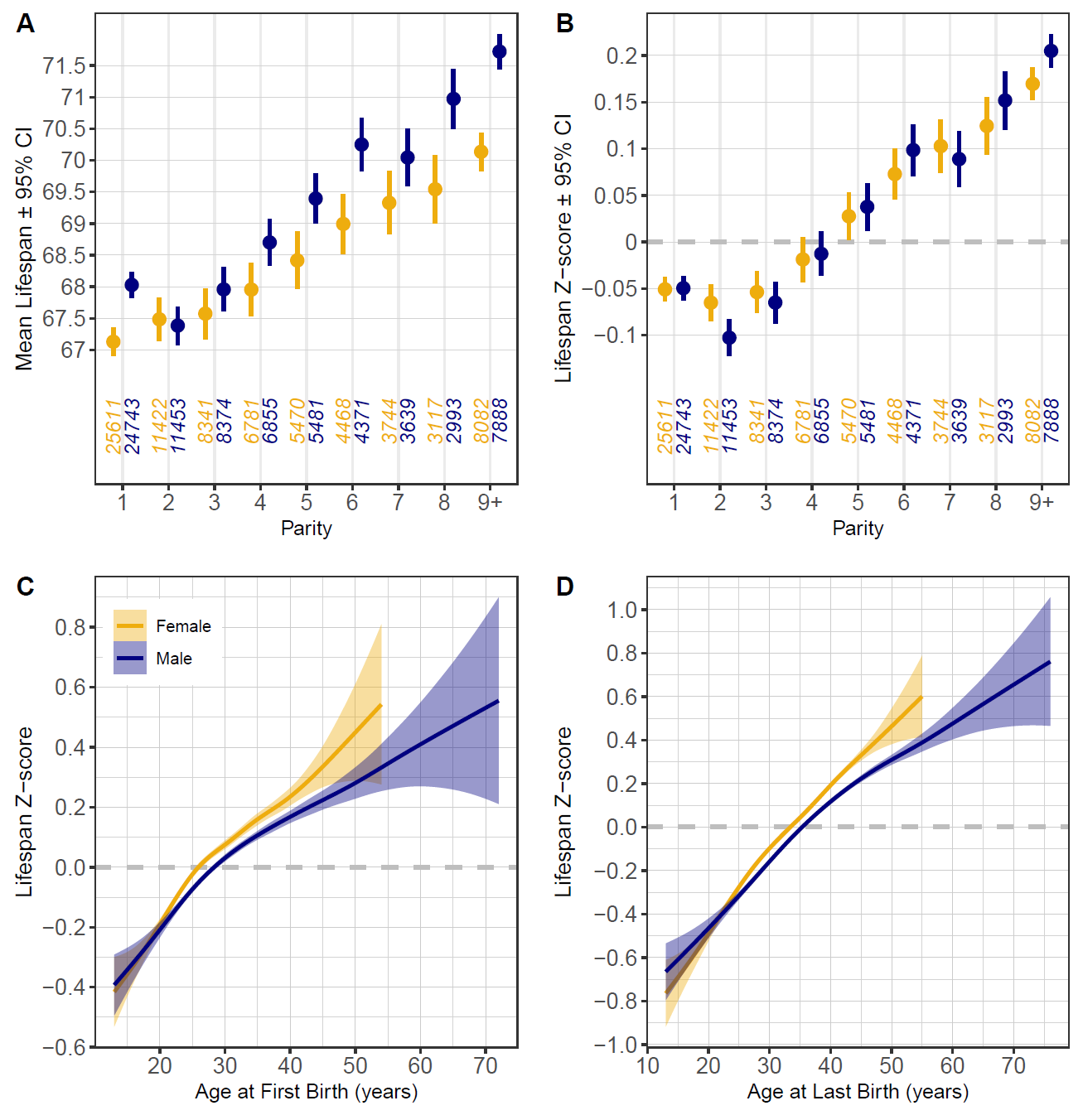

**Figure S6**. Associations between parity and crude lifespan (A) and lifespan z-scores, standardised within birth year and sex (B). Numbers at x-axis denote sample sizes per sex and parity. Birth-year adjusted predicted mean lifespans by parity category are shown in table S3 in the ESM. Associations between AFB (C) and ALB (D) vs lifespan z-scores, smoothed by GAM. Age is presented in years for the purpose of visualisation; age-group-specific linear slopes where both pairs of traits are adjusted for birth year are shown in the table S4 in ESM. Data for 13+ sample.

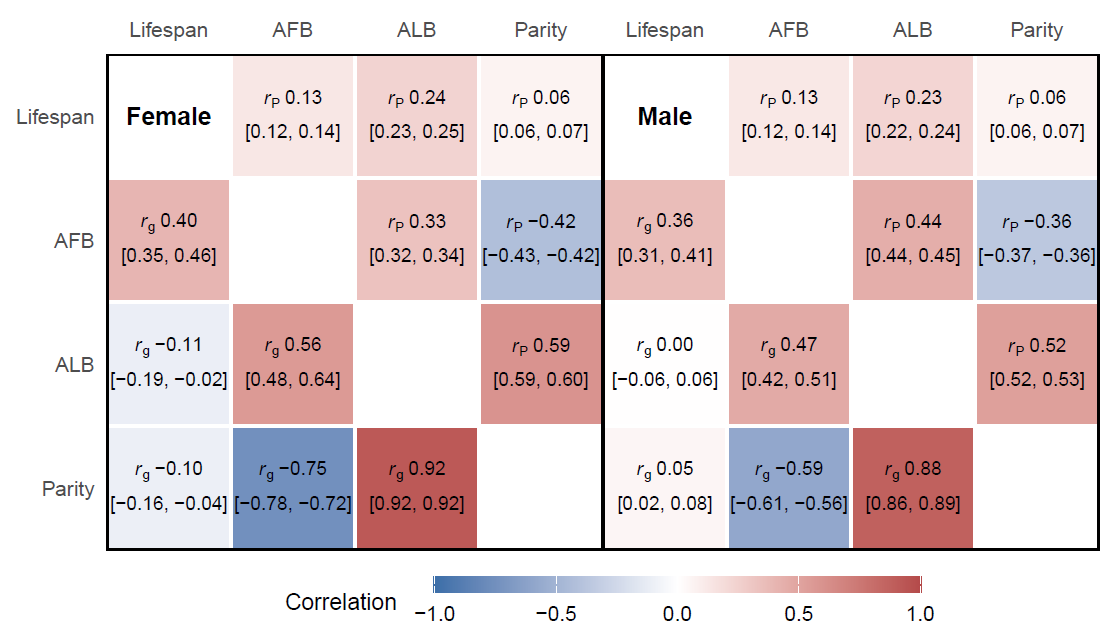

**Figure S7**. Phenotypic (above diagonal) and genetic (below diagonal) animal-model-derived correlations between lifespan and reproductive traits in 13+ sample. Numbers in parentheses denote 95% HPD intervals. Correlations with parity are converted to observed scale.

**Table S1.** Heritabilities for lifespan and reproductive traits among individuals who had reached age 13. Heritabilities are from single-sex univariate animal models; r_g_FM and r_p_FM are cross-sex genetic and phenotypic correlations from bivariate models in which the same trait in women and men was treated as two sex-limited responses. N phenot is the number of phenotyped individuals for each trait. N w. relatives is the number of phenotyped individuals having at least one phenotyped parent, offspring or sibling. Pairwise links counts relative pairs with r ≥ 0.125 (first cousins and closer); the model uses the complete pedigree, including more distant relationships. Gen. depth is average ancestral generation depth (maximum is 14). Narrow sense heritability estimates (h^2^) are presented as posterior means with the associated 95% Highest Posterior Density (HPD) intervals on the latent scale (h^2^_lat_) and, for parity also converted to the observed scale (h^2^_obs_) [35]. h^2^_obs_ was calculated at reference values of 3.0 children for females and 2.9 children for males. r_g_FM and r_p_FM are cross-sex genetic and phenotypic correlations between trait values.

| **Trait** | **Sex** | **h² (95% HPD)** | **N phenot.** | **N w. relatives** | **Pairwise links** | **Gen. depth** | **r_gFM (95% HPD) r_pFM (95% HPD)** |
| --- | --- | --- | --- | --- | --- | --- | --- |
| Lifespan | F | 0.23 (0.22-0.25) | 144 997 | 97 859 | 831 407 | 2.21 | 0.86 (0.81-0.91) |
|  | M | 0.23 (0.21-0.24) | 141 328 | 100 973 | 966 129 | 2.50 | 0.24 (0.23-0.25) |
| AFB | F | 0.34 (0.30-0.37) | 74 570 | 26 121 | 168 496 | 1.37 | 0.67 (0.52-0.88) |
|  | M | 0.49 (0.45-0.52) | 73 127 | 32 933 | 267 987 | 1.91 | 0.31 (0.28-0.33) |
| ALB | F | 0.13 (0.10-0.16) | 74 562 | 26 119 | 168 478 | 1.37 | 0.77 (0.57-0.95) |
|  | M | 0.34 (0.31-0.37) | 73 107 | 32 903 | 267 767 | 1.91 | 0.19 (0.17-0.21) |
| Parity lat | F | 0.27 (0.22-0.32) | 77 036 | 27 569 | 178 609 | 1.38 | 0.60 (0.49-0.70) |
|  | M | 0.73 (0.70-0.77) | 75 797 | 34 452 | 280 036 | 1.91 | 0.43 (0.39-0.47) |
| Parity obs | F | 0.15 (0.12-0.17) |  |  |  |  | 0.58 (0.48-0.70) |
|  | M | 0.43 (0.40-0.45) |  |  |  |  | 0.24 (0.22-0.26) |

**Table S2.** Phenotypic associations between lifespan and reproductive traits, by sex and surviving-age sample.

**Raw r**: Pearson correlation. **Partial r, YOB**: correlation between residuals after regressing both traits on ns(YOB, df = 5). **Partial r, YOB+region+urban**: both traits residualised jointly (in one model, same complete-case sample) on ns(YOB, df = 5) + province + urban/rural. **Regression b**: unstandardised slope of the first (outcome) trait on the second (predictor) trait from a single model adjusting jointly for ns(YOB, df = 5) + province + urban/rural; units are years of the outcome per year of the predictor (yr/yr), years of lifespan per additional child (yr/child), or additional children per year of the predictor (child/yr). AFB, age at first birth; ALB, age at last birth; YOB, year of birth. Partial correlation is the primary association measure; the regression coefficient is provided for its directional, unit-based interpretation.

| **Sex** | **Trait pair (outcome-predictor)** | **N** | **Raw r [95% CI]** | **Partial r, YOB [95% CI]** | **Partial r, YOB+region+urban [95% CI]** | **Regression b, YOB+region+urban [95% CI]** |
| --- | --- | --- | --- | --- | --- | --- |
| **45+ sample** | | | | | | |
| F | lifespan–children | 66 779 | -0.019 [-0.026, -0.011] | 0.005 [-0.002, 0.013] | 0.005 [-0.002, 0.013] | 0.019 [-0.009, 0.047] yr/child |
| F | lifespan–AFB | 64 598 | 0.034 [0.026, 0.041] | 0.037 [0.029, 0.044] | 0.039 [0.031, 0.046] | 0.087 [0.069, 0.104] yr/yr |
| F | lifespan–ALB | 64 617 | 0.019 [0.011, 0.027] | 0.040 [0.032, 0.048] | 0.042 [0.034, 0.050] | 0.078 [0.064, 0.093] yr/yr |
| F | AFB–ALB | 64 571 | 0.306 [0.299, 0.313] | 0.298 [0.291, 0.305] | 0.293 [0.286, 0.300] | 0.243 [0.237, 0.249] yr/yr |
| F | children–AFB | 64 598 | -0.397 [-0.403, -0.390] | -0.411 [-0.418, -0.405] | -0.420 [-0.426, -0.413] | -0.255 [-0.260, -0.251] child/yr |
| F | children–ALB | 64 617 | 0.598 [0.593, 0.603] | 0.589 [0.584, 0.594] | 0.587 [0.582, 0.592] | 0.296 [0.293, 0.299] child/yr |
| M | lifespan–children | 68 859 | 0.029 [0.021, 0.036] | 0.034 [0.027, 0.042] | 0.032 [0.025, 0.040] | 0.117 [0.090, 0.144] yr/child |
| M | lifespan–AFB | 66 408 | 0.023 [0.016, 0.031] | 0.024 [0.016, 0.031] | 0.022 [0.014, 0.029] | 0.043 [0.028, 0.058] yr/yr |
| M | lifespan–ALB | 66 434 | 0.048 [0.041, 0.056] | 0.052 [0.045, 0.060] | 0.049 [0.041, 0.057] | 0.079 [0.067, 0.092] yr/yr |
| M | AFB–ALB | 66 395 | 0.426 [0.420, 0.433] | 0.420 [0.414, 0.427] | 0.412 [0.406, 0.419] | 0.339 [0.334, 0.345] yr/yr |
| M | children–AFB | 66 408 | -0.333 [-0.340, -0.327] | -0.349 [-0.355, -0.342] | -0.360 [-0.366, -0.353] | -0.196 [-0.200, -0.193] child/yr |
| M | children–ALB | 66 434 | 0.559 [0.554, 0.564] | 0.548 [0.543, 0.553] | 0.545 [0.540, 0.551] | 0.245 [0.242, 0.248] child/yr |
| **13+ sample** | | | | | | |
| F | lifespan–children | 77 036 | 0.058 [0.051, 0.065] | 0.078 [0.071, 0.085] | 0.077 [0.070, 0.085] | 0.424 [0.385, 0.462] yr/child |
| F | lifespan–AFB | 74 570 | 0.110 [0.103, 0.117] | 0.114 [0.107, 0.121] | 0.114 [0.107, 0.121] | 0.375 [0.352, 0.398] yr/yr |
| F | lifespan–ALB | 74 562 | 0.188 [0.181, 0.195] | 0.209 [0.202, 0.216] | 0.211 [0.204, 0.218] | 0.560 [0.542, 0.579] yr/yr |
| F | AFB–ALB | 74 511 | 0.336 [0.330, 0.342] | 0.330 [0.324, 0.336] | 0.324 [0.318, 0.331] | 0.263 [0.258, 0.269] yr/yr |
| F | children–AFB | 74 570 | -0.375 [-0.381, -0.369] | -0.388 [-0.394, -0.382] | -0.397 [-0.403, -0.391] | -0.240 [-0.244, -0.236] child/yr |
| F | children–ALB | 74 562 | 0.598 [0.593, 0.602] | 0.589 [0.584, 0.594] | 0.587 [0.582, 0.591] | 0.288 [0.285, 0.291] child/yr |
| M | lifespan–children | 75 797 | 0.086 [0.079, 0.093] | 0.093 [0.086, 0.101] | 0.090 [0.083, 0.098] | 0.440 [0.405, 0.474] yr/child |
| M | lifespan–AFB | 73 127 | 0.100 [0.093, 0.107] | 0.101 [0.094, 0.109] | 0.098 [0.091, 0.106] | 0.258 [0.239, 0.277] yr/yr |
| M | lifespan–ALB | 73 107 | 0.184 [0.177, 0.191] | 0.192 [0.185, 0.199] | 0.189 [0.182, 0.196] | 0.400 [0.385, 0.415] yr/yr |
| M | AFB–ALB | 73 064 | 0.441 [0.436, 0.447] | 0.436 [0.430, 0.442] | 0.428 [0.422, 0.434] | 0.346 [0.341, 0.352] yr/yr |
| M | children–AFB | 73 127 | -0.317 [-0.324, -0.311] | -0.332 [-0.338, -0.325] | -0.343 [-0.349, -0.336] | -0.187 [-0.190, -0.183] child/yr |
| M | children–ALB | 73 107 | 0.564 [0.559, 0.569] | 0.553 [0.548, 0.558] | 0.551 [0.546, 0.556] | 0.243 [0.240, 0.246] child/yr |

**Table S3.** Birth-year-adjusted mean lifespan (predicted at the mean birth year) by parity category, from linear model: lm(lifespan ~ factor(parity) + ns(YOB, 5)).

| **Sex** | **Parity** | **N** | **Mean lifespan** | **95% CI** |
| --- | --- | --- | --- | --- |
| **45+ sample** | | | | |
| F | 1 | 21 830 | 71.22 | [70.97, 71.46] |
| F | 2 | 9 654 | 71.41 | [71.11, 71.71] |
| F | 3 | 7 058 | 71.43 | [71.10, 71.77] |
| F | 4 | 5 833 | 71.25 | [70.90, 71.61] |
| F | 5 | 4 744 | 71.62 | [71.24, 72.01] |
| F | 6+ | 17 660 | 71.50 | [71.24, 71.76] |
| M | 1 | 22 190 | 71.53 | [71.29, 71.77] |
| M | 2 | 10 071 | 71.35 | [71.06, 71.64] |
| M | 3 | 7 431 | 71.59 | [71.27, 71.90] |
| M | 4 | 6 206 | 71.68 | [71.33, 72.02] |
| M | 5 | 5 014 | 72.03 | [71.66, 72.40] |
| M | 6+ | 17 947 | 72.41 | [72.15, 72.66] |
| **13+ sample** | | | | |
| F | 1 | 25 611 | 65.51 | [65.18, 65.84] |
| F | 2 | 11 422 | 65.24 | [64.84, 65.64] |
| F | 3 | 8 341 | 65.47 | [65.03, 65.92] |
| F | 4 | 6 781 | 66.09 | [65.61, 66.57] |
| F | 5 | 5 470 | 66.93 | [66.41, 67.45] |
| F | 6+ | 19 411 | 68.69 | [68.33, 69.04] |
| M | 1 | 24 743 | 67.94 | [67.64, 68.23] |
| M | 2 | 11 453 | 67.06 | [66.70, 67.42] |
| M | 3 | 8 374 | 67.65 | [67.26, 68.05] |
| M | 4 | 6 855 | 68.49 | [68.06, 68.91] |
| M | 5 | 5 481 | 69.30 | [68.84, 69.77] |
| M | 6+ | 18 891 | 71.09 | [70.77, 71.41] |

**Table S4.** Within-age group linear slopes of regression of lifespan on age at first or last birth (years of lifespan per year of AFB or ALB), by reproductive-timing group, adjusted for birth year. Slopes are from a single linear model per sex and sample, lifespan ~ group + group-specific slope + ns(YOB, df = 5), i.e. birth year is adjusted for by including the spline as a covariate.

| **Sex** | **Trait** | **Age group** | **N** | **Slope** | **SE** | **p** |
| --- | --- | --- | --- | --- | --- | --- |
| **45+ sample** | | | | | | |
| F | AFB | <25 | 28 350 | 0.267 | 0.036 | <0.0000001 |
| F | AFB | 25-40 | 34 900 | 0.025 | 0.017 | 0.142 |
| F | AFB | >40 | 1 348 | 0.372 | 0.188 | 0.047 |
| M | AFB | <25 | 18 186 | 0.167 | 0.051 | 0.00098 |
| M | AFB | 25-40 | 44 531 | 0.028 | 0.015 | 0.052 |
| M | AFB | >40 | 3 691 | 0.126 | 0.049 | 0.010 |
| F | ALB | <25 | 7 204 | 0.401 | 0.076 | <0.0000001 |
| F | ALB | 25-40 | 44 075 | 0.052 | 0.013 | 0.0001 |
| F | ALB | >40 | 13 338 | 0.164 | 0.058 | 0.0045 |
| M | ALB | <25 | 3 931 | 0.203 | 0.112 | 0.0699 |
| M | ALB | 25-40 | 41 931 | 0.084 | 0.014 | <0.0000001 |
| M | ALB | >40 | 20 572 | 0.168 | 0.021 | <0.0000001 |
| **13+ sample** | | | | | | |
| F | AFB | <25 | 33 849 | 0.626 | 0.048 | <0.0000001 |
| F | AFB | 25-40 | 39 333 | 0.286 | 0.023 | <0.0000001 |
| F | AFB | >40 | 1 388 | 0.545 | 0.267 | 0.041 |
| M | AFB | <25 | 20 818 | 0.469 | 0.061 | <0.0000001 |
| M | AFB | 25-40 | 48 575 | 0.245 | 0.018 | <0.0000001 |
| M | AFB | >40 | 3 734 | 0.189 | 0.064 | 0.003 |
| F | ALB | <25 | 9 473 | 0.854 | 0.093 | <0.0000001 |
| F | ALB | 25-40 | 51 335 | 0.524 | 0.018 | <0.0000001 |
| F | ALB | >40 | 13 754 | 0.428 | 0.080 | <0.0000001 |
| M | ALB | <25 | 4 882 | 0.470 | 0.128 | 0.0003 |
| M | ALB | 25-40 | 47 259 | 0.465 | 0.017 | <0.0000001 |
| M | ALB | >40 | 20 966 | 0.287 | 0.027 | <0.0000001 |

**Table S5.** Per-model MCMC settings and convergence/efficiency diagnostics for the models whose estimates are reported in the manuscript. Single-sex univariate models provide the narrow-sense heritabilities of tables 1 and S1; cross-sex bivariate models provide the cross-sex genetic and phenotypic correlations (r_g FM, r_p FM) in those same tables and within-sex bivariate models provide the genetic and phenotypic correlations among traits in Figures 2 and S7. Chains were run with settings fixed a priori: a 30 000-iteration burn-in, no thinning, and one of two target run lengths. Long runs were executed as checkpointed batches concatenated into a single posterior sample, with the burn-in discarded once in the first batch and subsequent batches continuing the chain from the previous end state; because batches were of fixed size, realised chain lengths differ slightly from their nominal targets. Heidel pass fraction and Geweke pass fraction are the fractions of variance-covariance components passing the Heidelberger-Welch and Geweke (|z| < 1.96) tests, respectively. ESS: effective sample size; Nitt: total MCMC iterations; Stored: retained posterior samples.

| **Model** | **Family** | **Nitt** | **Stored** | **Min ESS** | **Median ESS** | **Heidel pass fraction** | **Geweke pass fraction** |
| --- | --- | --- | --- | --- | --- | --- | --- |
| mc_Female_AFB_YOBns5_motherUS_MF_conf_huge4 | gaussian | 1 280 000 | 1 250 000 | 749 | 2085 | 1.00 | 1.00 |
| mc_Female_AFB_YOBns5_motherUS_min45_MF_conf_huge4 | gaussian | 1 280 000 | 1 250 000 | 696 | 1863 | 1.00 | 0.67 |
| mc_Female_ALB_YOBns5_motherUS_MF_conf_huge4 | gaussian | 1 280 000 | 1 250 000 | 477 | 759 | 1.00 | 1.00 |
| mc_Female_ALB_YOBns5_motherUS_min45_MF_conf_huge4 | gaussian | 1 280 000 | 1 250 000 | 535 | 767 | 1.00 | 1.00 |
| mc_Female_children_YOBns5_motherUS_min45_pois_MF_conf_huge4 | poisson | 1 280 000 | 1 250 000 | 799 | 1980 | 1.00 | 0.00 |
| mc_Female_children_YOBns5_motherUS_pois_MF_conf_huge4 | poisson | 1 280 000 | 1 250 000 | 670 | 2098 | 1.00 | 0.33 |
| mc_Female_lifespan_YOBns5_motherUS_MF_conf_huge4 | gaussian | 1 280 000 | 1 250 000 | 1001 | 2589 | 1.00 | 0.67 |
| mc_Female_lifespan_YOBns5_motherUS_min45_MF_conf_huge4 | gaussian | 1 280 000 | 1 250 000 | 455 | 1833 | 1.00 | 0.67 |
| mc_Male_AFB_YOBns5_motherUS_MF_conf_huge4 | gaussian | 1 280 000 | 1 250 000 | 293 | 3026 | 1.00 | 0.33 |
| mc_Male_AFB_YOBns5_motherUS_min45_MF_conf_huge4 | gaussian | 1 280 000 | 1 250 000 | 232 | 2658 | 0.67 | 0.33 |
| mc_Male_ALB_YOBns5_motherUS_MF_conf_huge4 | gaussian | 1 280 000 | 1 250 000 | 558 | 2516 | 1.00 | 1.00 |
| mc_Male_ALB_YOBns5_motherUS_min45_MF_conf_huge4 | gaussian | 1 280 000 | 1 250 000 | 765 | 2344 | 1.00 | 0.67 |
| mc_Male_children_YOBns5_motherUS_min45_pois_MF_conf_huge4 | poisson | 1 280 000 | 1 250 000 | 1095 | 1692 | 1.00 | 1.00 |
| mc_Male_children_YOBns5_motherUS_pois_MF_conf_huge4 | poisson | 1 280 000 | 1 250 000 | 1102 | 1670 | 1.00 | 0.67 |
| mc_Male_lifespan_YOBns5_motherUS_MF_conf_huge4 | gaussian | 1 280 000 | 1 250 000 | 961 | 2153 | 1.00 | 1.00 |
| mc_Male_lifespan_YOBns5_motherUS_min45_MF_conf_huge4 | gaussian | 1 280 000 | 1 250 000 | 1344 | 1548 | 1.00 | 1.00 |
| mc_AFB_idh_YOBns5_motherUS_MF_conf_huge4 | gaussian+ gaussian | 1 280 000 | 1 250 000 | 359 | 1032 | 0.10 | 0.20 |
| mc_AFB_idh_YOBns5_motherUS_min45_MF_conf_huge4 | gaussian+ gaussian | 1 280 000 | 1 250 000 | 485 | 1251 | 1.00 | 1.00 |
| mc_ALB_idh_YOBns5_motherUS_MF_conf_lon4 | gaussian+ gaussian | 520 000 | 490 000 | 38 | 396 | 0.40 | 0.10 |
| mc_ALB_idh_YOBns5_motherUS_min45_MF_conf_lon4 | gaussian+ gaussian | 520 000 | 490 000 | 66 | 360 | 0.30 | 0.30 |
| mc_children_YOBns5_motherUS_min45_pois_MF_conf_huge4 | poisson+ poisson | 1 280 000 | 1 250 000 | 385 | 1503 | 1.00 | 0.70 |
| mc_children_YOBns5_motherUS_pois_MF_conf_lon4 | poisson+ poisson | 520 000 | 490 000 | 129 | 579 | 1.00 | 0.70 |
| mc_lifespan_idh_YOBns5_motherUS_MF_conf_huge4 | gaussian+ gaussian | 1 300 000 | 1 270 000 | 953 | 2754 | 0.60 | 0.10 |
| mc_lifespan_idh_YOBns5_motherUS_min45_MF_conf_huge4 | gaussian+ gaussian | 1 300 000 | 1 270 000 | 497 | 1603 | 0.80 | 0.80 |
| mc_Female_AFB_ALB_YOBns5_motherUS_MF_conf_lon4 | gaussian+ gaussian | 500 000 | 470 000 | 190 | 401 | 1.00 | 0.83 |
| mc_Female_AFB_ALB_YOBns5_motherUS_min45_MF_conf_lon4 | gaussian+ gaussian | 500 000 | 470 000 | 216 | 377 | 1.00 | 1.00 |
| mc_Female_children_AFB_YOBns5_motherUS_min45_pois_MF_conf_huge4 | poisson+ gaussian | 1 260 000 | 1 230 000 | 945 | 2385 | 1.00 | 0.30 |
| mc_Female_children_AFB_YOBns5_motherUS_pois_MF_conf_huge4 | poisson+ gaussian | 1 260 000 | 1 230 000 | 968 | 2795 | 1.00 | 1.00 |
| mc_Female_children_ALB_YOBns5_motherUS_min45_pois_MF_conf_lon4 | poisson+ gaussian | 500 000 | 470 000 | 1098 | 1980 | 1.00 | 0.70 |
| mc_Female_children_ALB_YOBns5_motherUS_pois_MF_conf_huge4 | poisson+ gaussian | 1 260 000 | 1 230 000 | 2702 | 4931 | 1.00 | 0.90 |
| mc_Female_children_lifespan_YOBns5_motherUS_min45_pois_MF_conf_huge4 | poisson+ gaussian | 1 260 000 | 1 230 000 | 408 | 1235 | 1.00 | 0.90 |
| mc_Female_children_lifespan_YOBns5_motherUS_pois_MF_conf_huge4 | poisson+ gaussian | 1 260 000 | 1 230 000 | 326 | 1723 | 0.90 | 0.20 |
| mc_Female_lifespan_AFB_YOBns5_motherUS_MF_conf_lon4 | gaussian+ gaussian | 500 000 | 470 000 | 155 | 605 | 1.00 | 0.58 |
| mc_Female_lifespan_AFB_YOBns5_motherUS_min45_MF_conf_lon4 | gaussian+ gaussian | 500 000 | 470 000 | 103 | 478 | 0.92 | 0.42 |
| mc_Female_lifespan_ALB_YOBns5_motherUS_MF_conf_lon4 | gaussian+ gaussian | 500 000 | 470 000 | 58 | 346 | 1.00 | 0.83 |
| mc_Female_lifespan_ALB_YOBns5_motherUS_min45_MF_conf_lon4 | gaussian+ gaussian | 500 000 | 470 000 | 102 | 273 | 1.00 | 0.92 |
| mc_Male_AFB_ALB_YOBns5_motherUS_MF_conf_lon4 | gaussian+ gaussian | 500 000 | 470 000 | 85 | 942 | 0.67 | 1.00 |
| mc_Male_AFB_ALB_YOBns5_motherUS_min45_MF_conf_lon4 | gaussian+ gaussian | 500 000 | 470 000 | 93 | 771 | 0.92 | 0.92 |
| mc_Male_children_AFB_YOBns5_motherUS_min45_pois_MF_conf_lon4 | poisson+ gaussian | 500 000 | 470 000 | 110 | 651 | 1.00 | 0.90 |
| mc_Male_children_AFB_YOBns5_motherUS_pois_MF_conf_huge4 | poisson+ gaussian | 1 260 000 | 1 230 000 | 254 | 1792 | 0.90 | 0.80 |
| mc_Male_children_ALB_YOBns5_motherUS_min45_pois_MF_conf_huge4 | poisson+ gaussian | 1 260 000 | 1 230 000 | 1117 | 2282 | 0.90 | 0.80 |
| mc_Male_children_ALB_YOBns5_motherUS_pois_MF_conf_huge4 | poisson+ gaussian | 1 260 000 | 1 230 000 | 742 | 2410 | 0.80 | 0.70 |
| mc_Male_children_lifespan_YOBns5_motherUS_min45_pois_MF_conf_lon4 | poisson+ gaussian | 500 000 | 470 000 | 265 | 459 | 1.00 | 0.90 |
| mc_Male_children_lifespan_YOBns5_motherUS_pois_MF_conf_huge4 | poisson+ gaussian | 1 260 000 | 1 230 000 | 513 | 1482 | 1.00 | 0.80 |
| mc_Male_lifespan_AFB_YOBns5_motherUS_MF_conf_lon4 | gaussian+ gaussian | 500 000 | 470 000 | 73 | 628 | 1.00 | 0.92 |
| mc_Male_lifespan_AFB_YOBns5_motherUS_min45_MF_conf_lon4 | gaussian+ gaussian | 500 000 | 470 000 | 69 | 645 | 1.00 | 0.92 |
| mc_Male_lifespan_ALB_YOBns5_motherUS_MF_conf_lon4 | gaussian+ gaussian | 500 000 | 470 000 | 200 | 611 | 1.00 | 0.50 |
| mc_Male_lifespan_ALB_YOBns5_motherUS_min45_MF_conf_lon4 | gaussian+ gaussian | 500 000 | 470 000 | 162 | 583 | 1.00 | 0.83 |

**Table S6.** Percentage of individuals giving birth after age 45 and age 50 (upper pane) and percentage of individuals having last birth after age 45 and 50 (lower pane).

| **Age at birth** | | | | |
| --- | --- | --- | --- | --- |
| **Sex** | **# of individuals** | **% > age 45** | **% > age 50** | **Median age at birth** |
| F | 368 397 | 0.441 | 0.036 | 30 |
| M | 349 494 | 4.061 | 0.953 | 32 |
| **Age at last birth** | | | | |
| **Sex** | **# of individuals** | **% > age 45** | **% > age 50** | **Median ALB** |
| F | 109 270 | 1.37 | 0.09 | 33 |
| M | 103 610 | 8.66 | 2.14 | 35 |

1.	de Villemereuil P., Schielzeth H., Nakagawa S., Morrissey M. 2016 General Methods for Evolutionary Quantitative Genetic Inference from Generalized Mixed Models. *Genetics* **204**(3), 1281-1294. (doi:10.1534/genetics.115.186536).
